# A genome-wide CRISPR screen reveals cancer-specific regulators of hyaluronan binding and cellular invasion

**DOI:** 10.64898/2026.02.06.704476

**Authors:** Jimmy Kim, Halen Kovacs, Anjali Parthasarathy, Shini Chen, Vignesh Krishnamoorthy, Mohammed Al-Seragi, May Zhao, Yu-Hsuan Huang, Pamela Austin Dean, Pauline Johnson, Calvin D. Roskelley, Simon Wisnovsky

**Author notes:** These authors contributed equally to this work (Jimmy Kim, Halen Kovacs, Anjali Parthasarathy, Shini Chen).

## Abstract

Metastatic spread of cancer cells is driven by binding between the cell-surface receptor CD44 and hyaluronan (HA) in the extracellular matrix. The specific oncogenes that drive increased CD44-HA binding in cancer remain poorly defined. Using a fluorescently labeled hyaluronan probe, we performed a genome-wide CRISPR screen to identify genes whose knockdown disrupts HA binding in breast cancer cells. We subsequently developed a bioinformatic analysis pipeline that enabled stratification and prioritization of cancer-specific regulators. This work provides a first-in-class resource for the identification of druggable targets to inhibit HA binding. We further validate RAB4A, a top hit from our screen, as a novel regulator of this process. Mechanistically, RAB4A KO dramatically reduces CD44 expression and inhibits the invasion of breast cancer cells through HA-rich matrices. This study validates a novel strategy for identifying regulators of cancer cell invasion and identifies immediate actionable targets for anti-metastatic therapy.

## Introduction

Tumor metastasis is a complex, multi-step process in which cells migrate from primary solid tumors and form secondary tumors at distant sites^1^. Metastasis causes most deaths from cancer, particularly for common solid malignancies like breast carcinoma^1^. There is thus an urgent need to characterize the molecular features that predispose cancer to metastasize and to identify specific targets for anti-metastasis therapy^1^. This metastatic process is facilitated by dynamic interactions between cancer cells and the extracellular matrix (ECM). Hyaluronan (HA), a key component of the ECM, is found in high abundance in many solid tumors^2-5^. HA is a linear polysaccharide glycosaminoglycan (GAG) that consists of repeating disaccharides of glucuronic acid and N-acetylglucosamine^5^. Its presence in the tumor microenvironment is associated with cancer progression^2^ and metastasis^2-4^. Functionally, adhesion of cancer cells to HA-rich ECM has been shown to facilitate extravasation and tissue homing of metastatic cancer^6-10^. HA interactions can also trigger pro-survival signaling pathways that enhance seeding and proliferation of metastatic cells^6-10^. HA binding thus drives metastasis in a range of different disease contexts.

In human cells, one significant HA-binding receptor is the cell-surface transmembrane glycoprotein CD44. Binding between CD44 and HA is tightly regulated by a complex, diverse set of molecular mechanisms. Various regulatory pathways can control: 1) transcription of the CD44 gene, 2) membrane trafficking and recycling of the CD44 protein, 3) alternative splicing of the CD44 mRNA and/or 4) post-translational modification of CD44^3,11^. In particular, variable glycosylation of the CD44 molecule with N-linked glycans^12^, sialic acid^13^ and/or chondroitin sulfate^14^ plays a critical role in regulating CD44’s ability to bind HA. In non-transformed cells, binding between CD44 and HA is often constitutively inhibited, even in cells that express CD44^10,15^. Cancer cells frequently remodel these regulatory pathways in ways that enhance their binding to HA, thus driving metastasis and tissue invasion.

There has been significant interest in inhibiting the CD44-HA interaction for anticancer therapy. Indeed, several therapeutic antibodies that neutralize CD44 binding to HA are currently in various stages of clinical development^16-18^. This broad-based therapeutic strategy, however, has the potential to elicit significant off-target effects^16-18^. For example, CD44-HA binding is critical for homing and tissue infiltration of activated immune cells^15^. CD44 is also a multi-functional molecule that can serve as a ligand for other important immune-regulatory receptors^19,20^. An alternative strategy would be to identify dysregulated pathways that drive elevated CD44-HA binding in cancer cells specifically. Such pathways, if discovered, would be prime targets for the development of more selective inhibitors. The complexity of CD44-HA regulation, however, has made it difficult to identify these types of molecules. A systematic strategy for identifying regulators of CD44-HA binding in cancer cells is thus urgently needed.

In this study, we developed a FACS-based chemical-genomic screening strategy to identify cellular factors that regulate the ability of cancer cells to bind HA. We applied this platform to studying MDA-MB-231 cells, a TNBC cell line that constitutively binds HA at high levels^21^. This work correctly identified known regulators of this process and defined a set of novel genes whose knockdown disrupts the CD44-HA interaction in cancer cells. Our novel method represents an efficient, modular approach to studying GAG-driven cellular interactions with the ECM. Validation of downstream hits further validated RAB4A as a key regulator of CD44-HA interactions and cellular invasion in a breast cancer cell line. These results are discussed in detail below.

## Results

### A genome-wide CRISPR screening platform uncovers regulators of CD44-HA binding in breast cancer cells

Building on previous work, we first developed a chemical probe to enable selective interrogation of HA-CD44 binding on intact cell surfaces. High molecular-weight HA (sourced from rooster comb, approximately 1000-4000 kDa by MW) was chemically conjugated with fluorescein (FL) using a previously established protocol (Fig. 1A)^22^. We then incubated this fluorescently labeled hyaluronan (FL-HA) with MDA-MB-231 breast cancer cells, which have been shown to express CD44 at high levels and to bind HA in the ECM^23^. Cell-surface binding of FL-HA was analyzed by flow cytometry. MDA-MB-231 cells bound strongly to FL-HA. Binding was greatly reduced by treatment with a CD44-blocking antibody. This data confirms prior work showing that CD44 is the major receptor for HA in this cell line, although other potential HA receptors (such as RHAMM) may also play a role (Fig. 1B)^24,25^. We then lentivirally transduced MDA-MB-231 cells with a dCas9KRAB construct, generating a chassis cell line that can be used for CRISPRi screening. KRAB is a transcriptional repression domain that mediates silencing of target genes^26^. Transduction of these cells with sgRNAs against CD44 led to strong silencing of both CD44 expression and FL-HA binding, confirming the efficiency of gene knockdown in our system (Fig. 1C-F).

**Figure 1.**
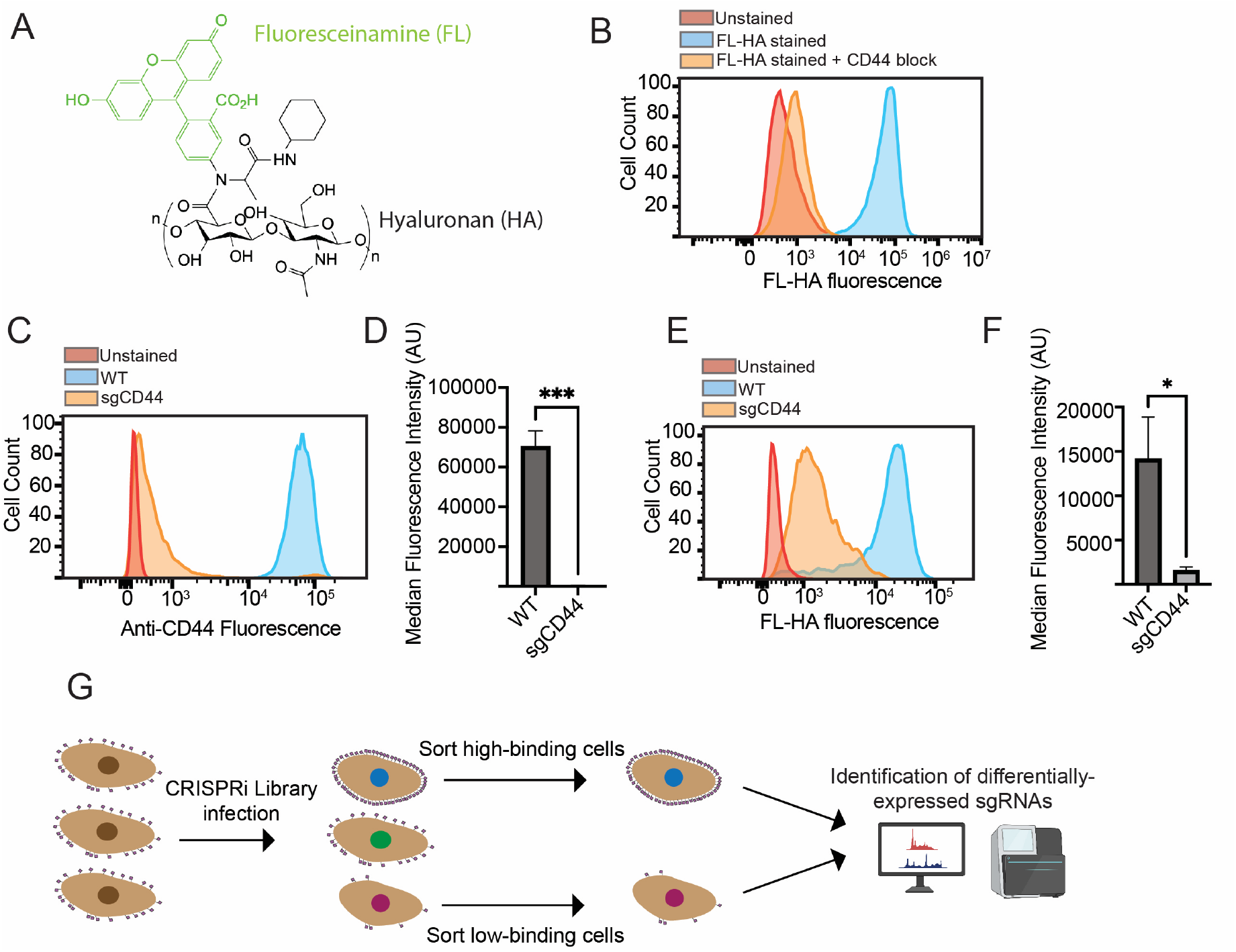
**A)** Chemical structure of Fluoresceinamine-Hyaluronan (FL-HA). **B**) MDA-MB-231 cells were incubated with FL-HA for 1 hour on ice at a 1:100 dilution and subsequently analyzed by flow cytometry. In some conditions, cells were pre-treated with a CD44 blocking antibody (HERMES-1 clone). For every 10,000 cells, 1 µL of blocking antibody was added to cells 30 minutes prior to incubation with FL-HA. **C**) MDA-MB-231-dCas9KRAB cells were transduced with an sgRNA against CD44. An antibody against CD44 (Hermes-3, 1:100) was then pre-complexed with an anti-mouse Alexa 488 secondary antibody (1.5 µg/mL) for 30 minutes and then incubated with the cells on ice. Cells were then analyzed by flow cytometry. A representative plot is shown. **D)** The median fluorescence intensity of CD44 staining is plotted for n=3 independent replicates. Statistical significance was determined using a Student’s two-tailed t-test, * indicates p<0.05, *** indicates p<0.001. **E)** MDA-MB-231-dCas9KRAB cells were transduced with an sgRNA against CD44 as in **1C**. Cells were then stained with FL-HA as in **1B**. A representative plot is shown. **F)** The median fluorescence intensity of FL-HA staining is plotted for n=3 independent replicates. Statistical significance was determined using a Student’s two-tailed t-test, * indicates p<0.05. **G)** Workflow of genome-wide CRISPRi screen. MDA-MB-231-dCas9KRAB cells were transduced with an sgRNA library containing 104,000 sgRNAs. Cells were subsequently stained with FL-HA as in **1B** and sorted by FACS to isolate low-staining (bottom 20%) and high-staining (top 20%) populations. sgRNAs were sequenced using an Illumina platform and sgRNA enrichment scores were calculated using MAGeCK.

We then proceeded to conduct a FACS-based CRISPR interference (CRISPRi) screen to identify genes whose knockdown altered CD44/HA binding. MDA-MB-231-dCas9KRAB cells were transduced with a genome-wide library of sgRNAs (∼18,500 genes, 5 sgRNAs/gene)^27^. Cells were then stained with FL-HA and sorted by FACS to isolate a “low-staining” (bottom 20% of the fluorescence distribution) and “high-staining” (top 20%) population (Fig. 1G). Next-generation sequencing was then performed to identify sgRNAs that were differentially enriched in either the low-staining or the high-staining population. Statistically significant hits were identified through MaGECK analysis^28^. As expected, the top hit of our CRISPRi screen was CD44, whose sgRNA-mediated knockdown significantly reduced staining with FL-HA (Fig. 2A). In addition, we identified 23 other hits whose knockdown significantly decreased FL-HA binding and 138 genes whose knockdown increased FL-HA binding (10% False Discovery Rate) (Supplementary Table 1). Henceforth, we will refer to hits whose knockdown decreased FL-HA binding as “positive regulators” and genes whose knockdown increased FL-HA binding as “negative regulators”.

**Figure 2.**
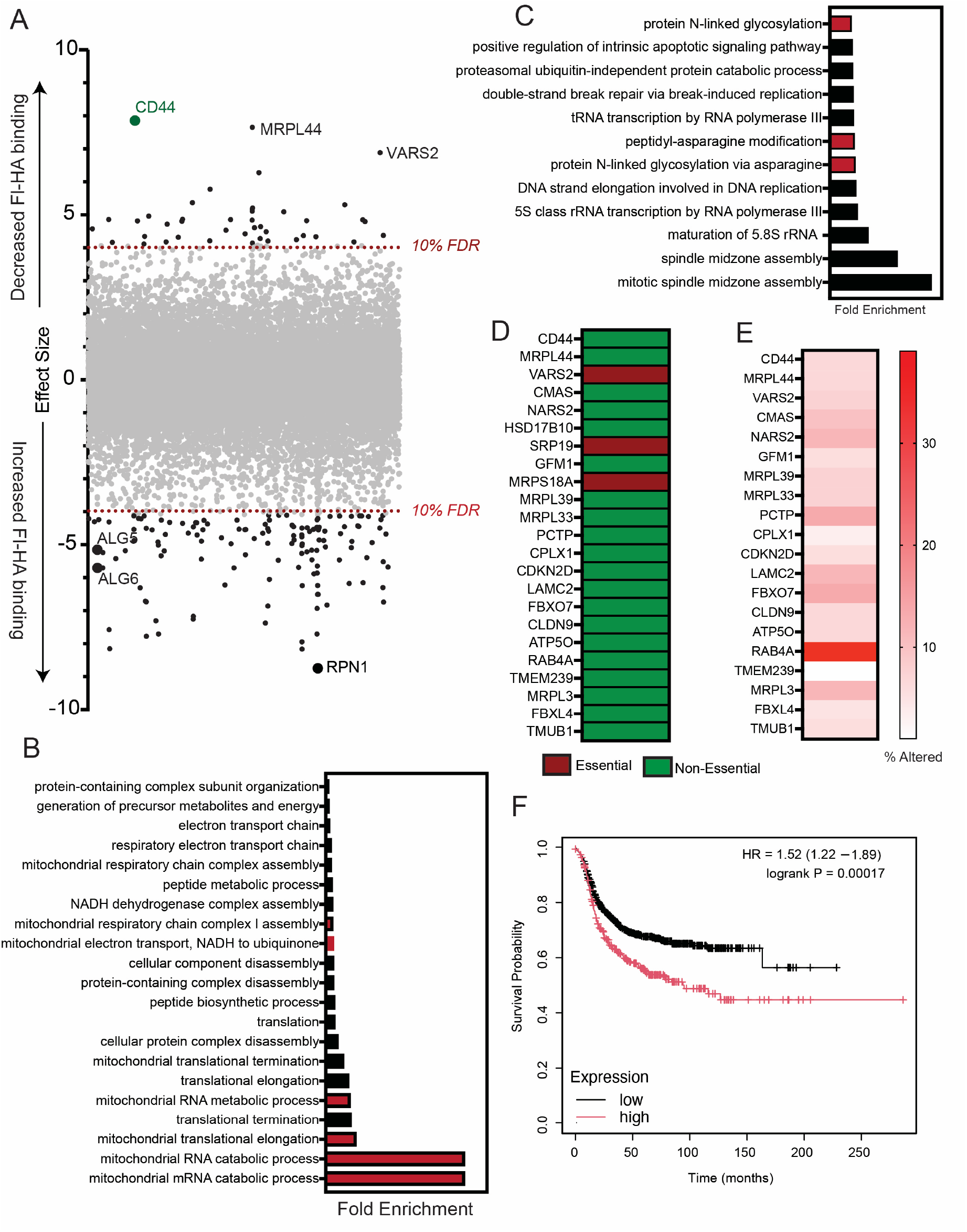
**A)** Results of genome-wide screen. An effect size was calculated and plotted for every gene as described in Materials & Methods. A positive effect size indicates gene knockdown reduced FL-HA binding, while a negative effect size indicates gene knockdown increased binding. False discovery rate (FDR) cut-offs are indicated. **B)** Gene Ontology (GO) enrichment analysis of gene hits with a positive effect score was performed using GOrilla. The top 20 terms by fold enrichment are ranked. All enrichments were statistically significant with an FDR (q-value) below 0.01. GO terms associated with mitochondrial translation were commonly found and are highlighted in red. **C)** Analysis of genes with a negative effect score was performed as in **1B**. GO terms associated with glycosylation were commonly found and are highlighted in red. **D)** Gene hits with a positive effect score were analyzed in DepMap to identify essential genes. Genes that have been found to be important for cell growth are highlighted in red. **E)** Gene hits with a positive effect score were analyzed in TCGA as described in Material & Methods. The percentage of patient samples showing dysregulation in the given gene (copy number or mRNA amplification) is plotted. **F)** Survival analysis for breast cancer patients with low and high expression of RAB4A was performed as described in Materials & Methods. HR = Hazard ratio, 95% confidence intervals plotted.

We next performed gene ontology (GO) enrichment analysis to identify specific pathways and molecular functions that were associated with these top hit genes. For positive regulators, this analysis revealed a strong enrichment for GO terms related to mitochondrial oxidative phosphorylation (Fig. 2B). Genes encoding components of the mitochondrial ribosomal machinery (e.g., MRPL44, MRPS18A) and mitochondrial tRNA synthetases (e.g, VARS2, NARS2) were particularly highly ranked positive regulators. The mechanistic basis for this effect is not clear, but several explanations are possible. Alterations in mitochondrial function can impact nuclear gene expression by modulating levels of acetyl-CoA, an essential cofactor for histone acetylation^29^. Global changes in cellular metabolic balance may also indirectly affect CD44 glycosylation, which is known to regulate HA binding^14^. Regardless of the underlying mechanism, our data point to a previously undiscovered connection between mitochondrial metabolism and acquisition of an HA-binding phenotype. Given the extensive metabolic reprogramming that occurs during breast cancer progression^30^, this finding may have important implications for understanding regulation of the metastatic process.

Amongst negative regulators, we observed a strong enrichment for GO terms associated with N-linked glycosylation (Fig. 2C). Specific hits included several genes (ALG5, ALG6, RPN1) involved in biosynthesis of the dolichol-linked oligosaccharide that serves as a donor for protein glycosylation^31^. This result further validates our screening dataset, as N-glycosylation of CD44 has previously been found to negatively regulate binding between CD44 and HA^32,33^. The gene CMAS, which is a key enzyme involved in the biosynthesis of sialic acid, was also recovered as a hit. This finding similarly aligns with prior work showing that sialylation of N-linked glycans can alter the capacity of CD44 to bind HA^13^. Other GO terms of interest included some related to mitosis and cytokinesis. Aurora Kinase B (AURKB), a critical regulator of this process, was one of the top positive regulators. Our screen thus broadly annotates a range of new biological pathways that drive CD44-HA binding in cancer cells. We present our full results in Supplementary Table 1 as a general resource for further investigation of these novel mechanistic connections.

### Pan-cancer genomic analysis reveals cancer-specific drivers of CD44-HA binding

We next sought to leverage our dataset to identify factors that drive CD44-HA binding and whose activity might be specifically dysregulated in cancer cells. We focused this analysis on genes whose knockdown reduced HA binding, as these genes are likely to be of greater interest as therapeutic targets for cancer metastasis. First, we consulted DepMap to identify genes that had been found to be broadly essential for cell growth across many cell line models. We reasoned that genetic knockout of such genes may produce pleiotropic effects on cell function, making it difficult to define specific mechanistic connections to CD44-HA biology. Gene essentiality can also make it challenging to establish stable knockout cell lines for downstream validation experiments. Several genes were excluded from further analysis on this basis (Fig. 2D). These notably included many of the mitochondrial translation genes we had previously noted above (Fig. 2D). Gene hits that were not annotated as commonly essential were carried forward into further analysis.

To more systematically prioritize hits, we next looked for genes that showed evidence of genetic dysregulation in breast cancer patient samples. We analyzed DNA and RNA sequencing data from a published cohort of 818 patients with invasive breast cancer^34^. From this dataset, we quantitated the percentage of patients that showed structural alterations in copy number (gene amplification) and/or elevated mRNA expression for each of our genes of interest. Several genes from our list were significantly upregulated (either in copy number, mRNA expression, or both) in metastatic breast cancer tissue compared to normal diploid cells. This analysis is presented in Fig. 2E. A single gene showed significantly more evidence of genetic dysregulation than any other on the list - the RAB GTPase RAB4A. Copy number amplification and/or mRNA upregulation of RAB4A was observed in nearly 40% of all breast cancer invasive carcinomas, implying a potentially broad significance for this gene in oncogenesis. On average, RAB4A mRNA expression was highest in metastatic tumor samples when compared to primary tumors and healthy tissue (Supplementary Fig. 1). Finally, in aggressive basal-like cancers, high RAB4A expression was also associated with significantly worse patient survival (Fig. 2F). These strong clinical correlations motivated us to focus on RAB4A as a key target gene of interest.

RAB4A is a Ras-family GTPase that localizes to the endo-lysosomal system^35^. Genetic depletion of RAB4A has been shown to affect the recycling of cell-surface proteins from the early endosome to the cell surface^35,36^. Impaired RAB4A function can cause mis-localization of proteins to the lysosome and other organelles^35,36^. In breast cancer, genetic knockdown of RAB4A has been found to impede tumor formation and metastasis *in vivo*^37,38^. The mechanistic basis for this effect is unclear: various studies have implicated a role for RAB4A in epithelial-mesenchymal transition^37^, maintenance of cancer stemness^38^, and cellular invasion through modulation of integrin recycling^39^. However, a specific connection between RAB4A, CD44 and hyaluronan binding has never been made. We thus wondered whether RAB4A may drive cancer metastasis by up-regulating binding between CD44 and HA.

### CRISPR-Cas9 KO of RAB4A significantly ablates CD44 expression and FL-HA binding

First, we lentivirally transduced MDA-MB-231 cells with a plasmid encoding Cas9 and a sgRNA targeting RAB4A. In parallel, cells were also transduced with a non-targeting sgRNA as a control. We subsequently used ICE (Synthego) to quantify indel formation at the RAB4A locus targeted by our sgRNA^40^. This analysis revealed a mixed polyclonal population of cells mainly containing a +1 insertion in the RAB4A gene, which would be expected to produce a nonsense mutation (Fig 3A). Overall editing efficiency was over 95% by this metric. In parallel, we also transduced MDA-MB-231-dCas9KRAB cells with an sgRNA targeted at the RAB4A promoter region. Near-complete silencing of RAB4A gene expression was observed using RT-qPCR (Supplementary Fig. 2). We also observed a complete ablation of RAB4A expression at the protein level in both cell lines using Western blotting (Fig. 3B, Supplementary Fig. 3). Below, we will refer to the CRISPR-edited cell line as “RAB4A KO cells”, while referring to the CRISPRi cell line as “RAB4A KD cells”.

**Figure 3.**
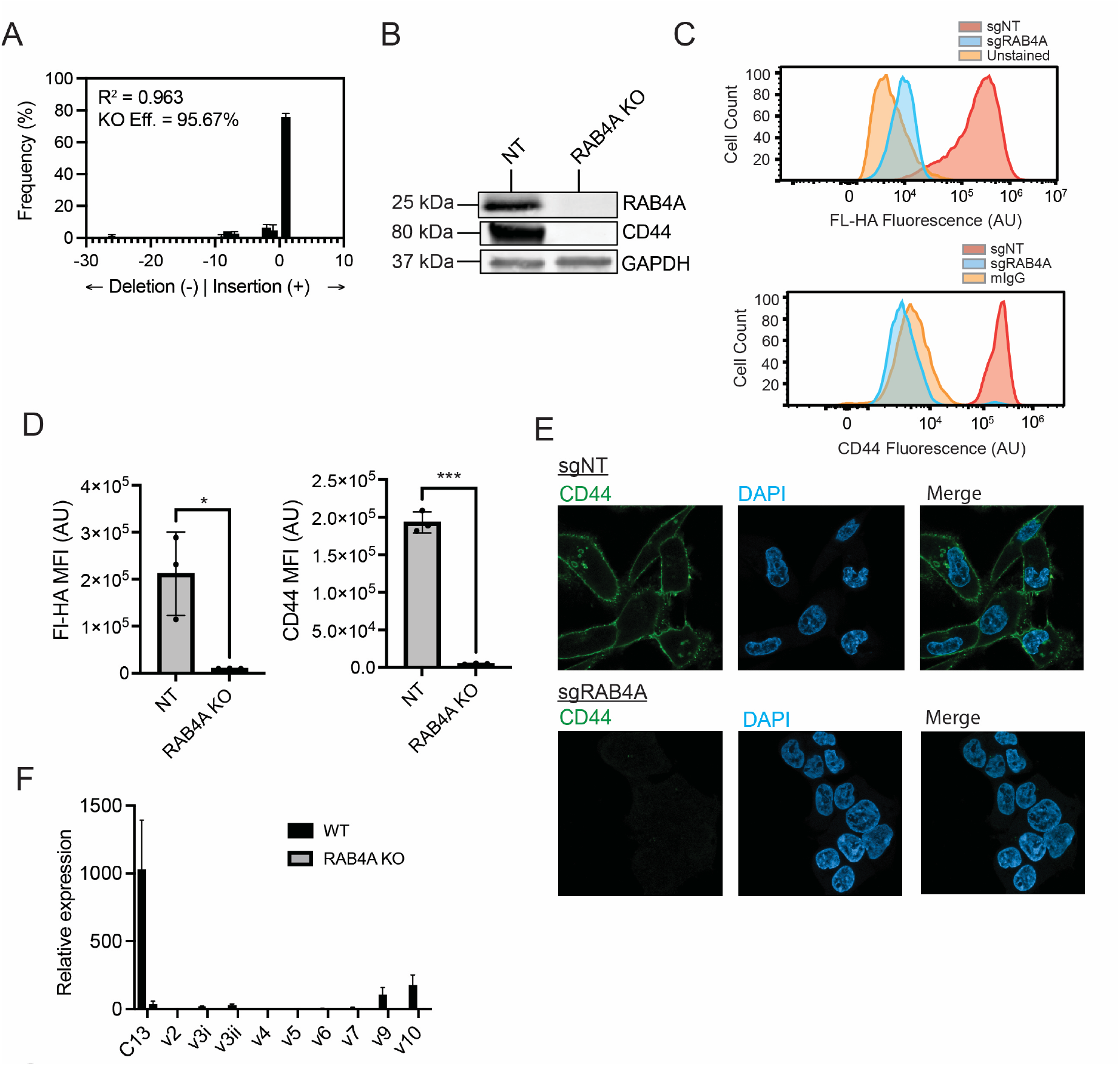
**A)** MDA-MB-231 cells were lentivirally transduced with Cas9 and either a non-targeting (NT) sgRNA or an sgRNA targeting RAB4A. A genomic region surrounding the target cut site was then amplified by PCR. Sanger sequencing was performed to determine the extent of indel formation at this locus. **B)** sgNT and sgRAB4A cells were lysed using RIPA buffer. An immunoblot was performed using antibodies targeting RAB4A, CD44 and GAPDH. **C)** sgNT and sgRAB4A cells were incubated with FL-HA and anti-CD44 antibodies and analyzed by flow cytometry as in **1C-F**. A representative plot is shown. **D)** The median fluorescence intensity (MFI) of FL-HA and CD44 staining in sgNT and sgRAB4A cells is plotted for n=3 biological replicates. **E)** sgNT and sgRAB4A cells were seeded overnight on imaging plates and fixed using 4% PFA. Following blocking and permeabilization, cells were incubated with an antibody against CD44 at a 1:500 dilution overnight, followed by a goat-anti-mouse Alexa 488 at 1:150. **F)** RNA was extracted from sgNT and sgRAB4A cells and CD44 mRNA expression levels were measured by qPCR using primers specific for different CD44 splice variants.

To confirm the initial result from our screen, we next stained sgNT, RAB4A KO and RAB4A KD cells with FL-HA. Loss of RAB4A produced a robust decrease in FL-HA staining, thus validating RAB4A as a regulator of CD44/HA binding (Fig. 3C-D, Supplementary Fig. 4). Both RAB4A KO and RAB4A KD cells displayed an identical phenotype. As these cell lines were generated through orthogonal CRISPR-based approaches, this result strongly suggests that loss of FL-HA binding is directly linked to loss of RAB4A, rather than an off-target effect of gene manipulation. Remarkably, the overall effect of RAB4A KO/KD was similar in magnitude to that produced by direct KD of the CD44 gene (Fig. 1C). We therefore wondered whether RAB4A may regulate the expression of CD44 at the cell surface. Indeed, flow cytometry analysis showed that RAB4A KO/KD cells exhibited significantly lower levels of staining with an anti-CD44 antibody (Fig. 3C-D, Supplementary Fig. 5). Analysis of cells by immunofluorescence microscopy showed a similar loss in cell-surface CD44 expression (Fig. 3E).

**Figure 4.**
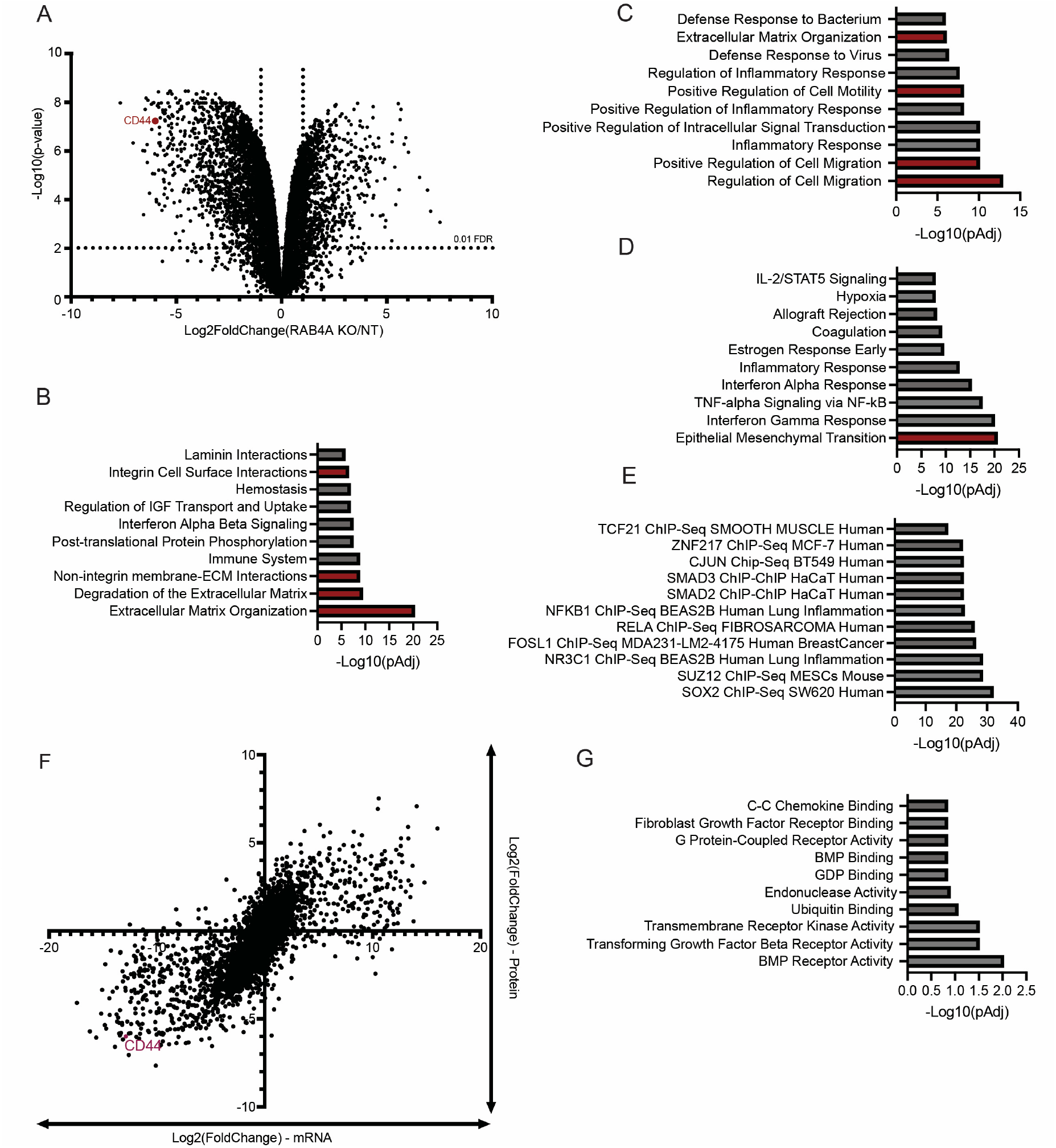
**A)** sgNT and sgRAB4A cells were analyzed by mass spectrometry proteomics. The volcano plot indicates relative changes in protein abundance upon RAB4A KO. A false discovery rate (FDR) threshold of 0.01 is indicated. **B)** Reactome pathway analysis of genes depleted in RAB4A KO cells is shown. **C)** Gene Ontology (GO) enrichment analysis of genes depleted in RAB4A KO cells is shown. **D)** MSigDB Hallmarks analysis of genes depleted in RAB4A KO cells is shown. For **B-D**, functional terms associated with cellular migration are highlighted in red. **E)** Transcription factor analysis of genes depleted in RAB4A KO cells is shown. **F)** Plot indicates the differential expression of genes at the mRNA (transcriptomic) and protein (proteomic) level following RAB4A KO. **G)** GO enrichment analysis of genes depleted at the protein level but not the mRNA level in RAB4A KO cells. All bioinformatic analysis was conducted as described in Materials & Methods.

**Figure 5.**
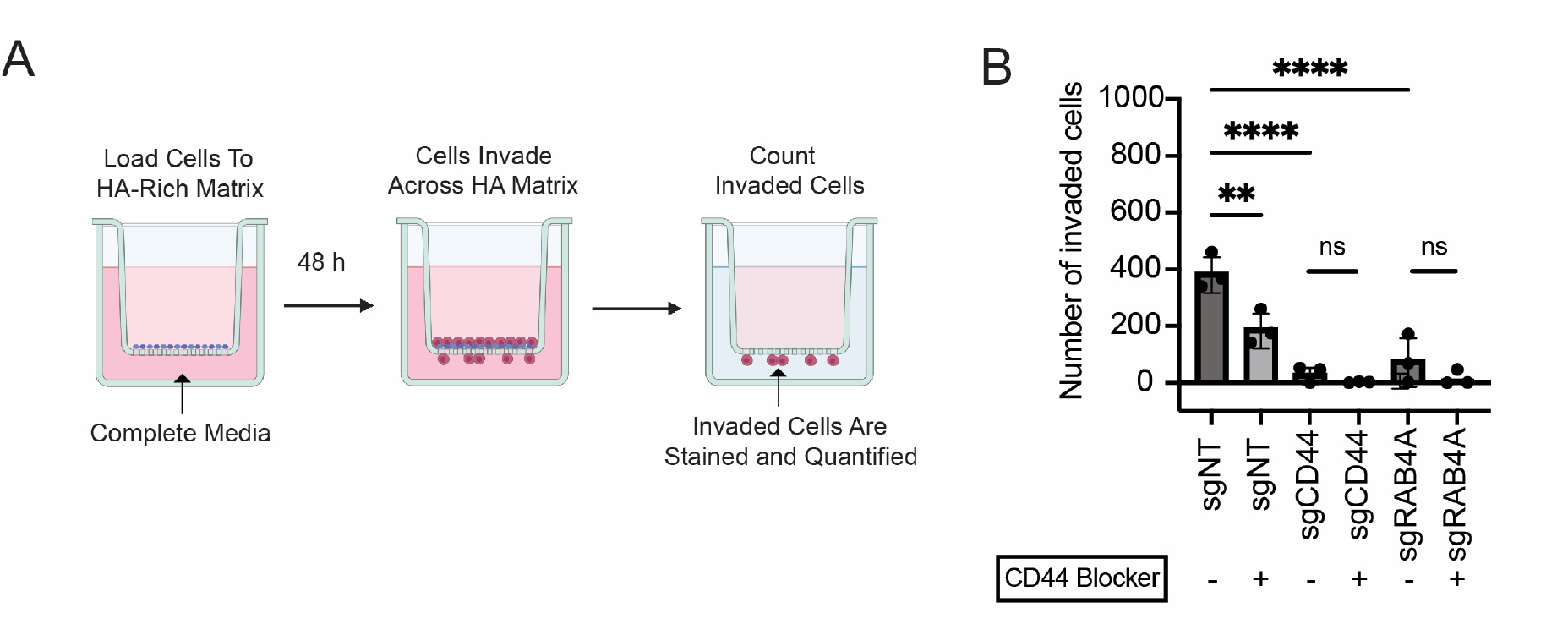
**A)** Workflow of transwell invasion assay. sgNT, sgCD44 and sgRAB4A cells were seeded on top of transwell inserts that had been coated with high MW HA. Cells were then allowed to invade for 48 h. Cells entering the matrix were counted using crystal violet staining and microscopy. **B)** The average number of invading cells is plotted for n=3 biological replicates. P-values were determined using an ordinary One-Way ANOVA. * indicates p<0.05, ** indicates p<0.01, *** indicates p<0.001, **** indicates p<0.0001.

### RAB4A KO initiates large-scale reprogramming of breast cancer cells towards a non-invasive cellular phenotype

We therefore interrogated how RAB4A regulates CD44 cell-surface residency. A set of prior studies have shown that RAB4A drives epithelial-mesenchymal transition (EMT) in several cancer cell lines^37,43^. As RAB4A is primarily involved in endosomal recycling of cell-surface proteins, the underlying molecular mechanism for this process is not clear^37,43^. However, there is some evidence that RAB4A alters a downstream signaling pathway involving the RAC1 GTPase as well as transcription factors known to drive EMT^37,41^. We hypothesized that this transcriptional reprogramming may affect the expression of CD44. Indeed, RAB4A KO cells exhibited a near-complete ablation of CD44 protein expression when measured by immunoblotting (Fig. 3B). We also tested the expression of CD44-encoding mRNAs by RT-qPCR and found that RAB4A KO produced a general collapse in CD44 mRNA levels (Fig. 3F). Notably, we measured a general reduction in mRNA expression for all known splice variants of CD44, indicating that RAB4A KO likely affects transcription of the CD44 gene rather than stability of specific CD44 mRNAs (Fig. 3F). Interestingly, analysis of breast cancer patient samples (TCGA) also showed a moderate (PCC=0.33) correlation between RAB4A and CD44 mRNA expression (Supplementary Fig. 6). Taken together, these data strongly suggest that RAB4A regulates CD44 gene expression, thus impeding the ability of cancer cells to bind HA.

Given these effects on gene transcription, we next chose to more broadly investigate the cellular effects of RAB4A KO. We lysed sgNT and sgRAB4A-transduced cells and subjected lysates to bottom-up proteomics analysis using mass spectrometry. This experiment demonstrated that RAB4A KO induces broad and dramatic remodeling of protein expression patterns in MDA-MB-231 cells (Fig. 4A). Out of 8863 proteins identified, 1180 showed significant (effect size > 2, FDR < 0.01) changes in abundance following RAB4A KO. CD44 was one of the most differentially expressed proteins in this dataset, confirming our previous results. REACTOME and GO term analysis revealed that RAB4A KO broadly alters expression of proteins involved in extracellular matrix organization (Fig. 4B), regulation of cell migration (Fig. 4C) and epithelial mesenchymal transition (Fig. 4D). Transcription factor analysis showed SOX2, a known regulator of EMT, as a possible upstream regulator of these changes in gene expression (Fig. 4E)^42^. These data strongly suggest that RAB4A KO perturbs multiple, fundamental signalling pathways that regulate the metastatic phenotype.

Our prior results suggested that RAB4A KO affects the abundance of CD44 mRNA. We therefore hypothesized that the majority of the proteomic changes we observed likely stem from changes in gene transcription. To assess this idea, we subjected sgNT and sgRAB4A cells to whole transcriptome analysis by RNA-seq. As expected, this study showed a dramatic remodeling of cellular mRNA expression, similar to what we observed at the protein level (Supplementary Fig. 6). Once again, CD44 was detected as one of the top differentially expressed genes in sgRAB4A cells. These data clearly indicate that disruption of RAB4A induces remarkable remodeling of gene expression patterns in breast cancer cells. Overall, RAB4A knockout seems to redirect cells towards a less invasive and migratory phenotype. CD44 is one of the primary proteins affected by this reprogramming, but by no means the only one. Notably, we would highlight that these findings align with several other studies of RAB4A’s function in breast cancer^37-39^.

These findings raise an interesting mechanistic question. RAB4A is an endosomal GTPase whose primary role is to drive fast recycling of cell membrane proteins from early endosomes to the cell surface. While RAB4A has previously been suggested to affect signaling through RAC1 and/or NOTCH, it is not clear how RAB4A KO would directly affect such signaling pathways^43,44^. RAB4A is not an essential gene for cellular fitness. It is also just one of many RAB proteins that perform similar functions in the endo-lysosomal system. It is thus quite unexpected that knockout of this one gene would produce such a striking phenotype. However, one possible mechanism is the effect that RAB4A KO may have on the localization of cell-surface signaling receptors. Constitutive signaling through a variety of cell membrane receptors is required for maintaining breast cancer cells in an invasive, mesenchymal state^43,44^. RAB4A may be required for promoting recycling and cell-surface localization of such receptors. In this model, RAB4A KO may induce aberrant localization of such proteins to the lysosome and subsequent proteolytic degradation. Loss of signaling potential could then initiate the large-scale rewiring of cellular gene expression patterns that we observe in our KO cells.

We explored this idea by integrated analysis of our proteomic and transcriptomic datasets. In general, mRNA and protein expression values in these data displayed a remarkable correlation. This pattern indicates that the vast majority of differentially expressed proteins in RAB4A KO cells stem from secondary effects of RAB4A KO on cell signaling, rather than resulting from immediate effects of RAB4A KO on endosomal recycling (Fig. 4F). However, we were able to identify a subset of proteins that exhibited a dramatic reduction in expression at the protein level, but not the mRNA level. We reasoned that some of these proteins might be directly perturbed by the immediate effects of RAB4A KO on endolysosomal trafficking. Interestingly, GO term analysis revealed a strong functional enrichment for proteins with transmembrane receptor signaling activity (Fig. 4G). The bone morphogenic protein (BMP) receptor family were particularly well-represented, with both BMPR2 and BMPR1A showing strong reduction at the protein but not the mRNA level. Interestingly, BMPR2 signaling has previously been shown to drive EMT and CD44 expression in MDA-MB-231 cells^45^. Several SMAD proteins, which are downstream effectors of BMPR signaling, were also recovered as possible transcriptional regulators that could underlie the gene expression changes observed in our data (Fig. 4E). These results are not definitive, and it is likely that RAB4A KO alters cell signaling through multiple parallel mechanisms. However, they do suggest that loss of RAB4A may induce post-translational destabilization of specific cell surface receptors, thus re-wiring cell signaling pathways within breast cancer cells.

### RAB4A KO reduces invasiveness of breast cancer cells *in vitro*

CD44-HA binding is essential for cancer cell invasion and migration through HA-rich ECM. We thus finally tested whether RAB4A KO would impede the ability of cancer cells to invade HA-rich matrices. We coated transwell inserts with high-molecular weight HA to form a solidified HA-containing matrix. MDA-MB-231 cells were then seeded on top of this matrix. After allowing cells to invade the HA layer, uninvaded cells on the top of the matrix were scraped off and invading cells were counted by Crystal Violet staining (Fig. 5A). WT MDA-MB-231 cells exhibited substantial invasion through HA. Invasion of WT cells was partially blocked by addition of a CD44-blocking antibody. RAB4A KO cells, conversely, displayed a dramatic reduction in invasion. Addition of a CD44-blocking antibody to RAB4A KO cells had no further effect on invasion. The impact of RAB4A KO on invasion was also similar to that produced by direct knockdown of CD44 (Fig. 5B). These data thus confirm that RAB4A drives invasion of breast cancer cells through HA-rich ECM by regulating expression of the HA receptor CD44.

## Discussion

In this study, we conducted a genome-wide CRISPR screen to map regulators of CD44-HA binding in breast cancer cells. To our knowledge, this is the first unbiased functional genomic study that has interrogated regulation of this critical biological process. Our CRISPR screen results provide an important resource for researchers interested in probing the mechanisms that underlie invasion and metastasis of cancer cells. Systematic triage for non-essential oncogenes singled out the small GTPase RAB4A as the top hit. Other genes on our list may similarly regulate CD44-HA binding and could also be targets of interest in different cancer types and genetic backgrounds.

These studies add to a body of literature that has implicated RAB4A as a driver of cell invasion and metastasis in breast cancer^37,39^. RAB4A mRNA expression is also clearly upregulated in a wide range of invasive breast cancer patient samples. When considered together, all these data frame RAB4A as a “master regulator” that controls cancer cell invasiveness by regulating key cell surface adhesion molecules. Additionally, RAB4A -/-mice are viable and exhibit no obvious phenotypes other than mild changes in creatine metabolism (IMPC, 2025. *Rab4a phenotype data set*, Data release vM15. MGI reference J:211773). In theory, inhibition of RAB4A could thus allow for selective blockade of CD44-HA binding in cancer cells without deleterious effects on normal tissues. While the RAB GTPase family has previously been thought to be undruggable, this view has been rapidly changing in recent years. One recent study reported that these enzymes possess a conserved, cryptic allosteric pocket that can be effectively liganded with small molecules^46^. Indirect inhibition of RAB prenylation^47^ and carboxylmethylation^39^ has also been explored as a potential strategy for blockade of RAB-family enzymes. These avenues for targeting the RAB4A-CD44-HA axis will be interesting to explore in future work.

Our work provides some basic insight into the mechanism by which RAB4A regulates CD44-HA binding. However, the details of how RAB4A selectively controls transcription of CD44 have not been fully elucidated. Different RAB-family proteins exhibit differing membrane localization within the endo-lysosomal system and interact with a distinct set of cargo proteins^48,49^. The molecular basis for this cargo specificity is not yet clear. Notably, RAB4A was the only RAB protein to emerge as a hit in our screen, indicating a specific importance for this protein rather than RAB-mediated recycling in general. Our data suggest that RAB4A may induce transcriptional reprogramming in cancer cells by altering cell surface residency of key cell surface signaling proteins like BMPRs. Changes in stability of these unknown proteins might then drive downstream changes in cell signaling and transcriptional reprogramming. Future work could focus on confirming and defining these interactions through protein truncation and/or split proximity labeling assays. It would also be interesting to determine how RAB4A KO affects cell surface residency of specific known oncogenic signaling proteins. Cell-surface capture proteomics could be usefully applied towards answering such questions.

One major limitation of our study is that our work was conducted in a single breast cancer cell line (MDA-MB-231 cells). We did profile a number of other TNBC cell lines (e.g, BT-549, MDA-MB-468 and HS578T cells) to identify other models that may be suitable for studying regulation of CD44-HA binding. Unfortunately, in our hands, BT-549 and HS578T cells displayed no appreciable binding to FL-HA (Supplementary Fig. 7A-B). MDA-MB-468 cells did bind FL-HA, but did not express CD44, indicating that hyaluronan binding is likely mediated by alternate receptors like RHAMM in this cell line (Supplementary Fig. 7C). We would note that our findings align well with prior literature, which has also largely used MDA-MB-231 cells as the primary model for studying HA binding in cancer^25,50-52^. Like many cancer phenotypes, it is likely that HA binding may often be lost following removal of cancer cells from their native microenvironment and adaptation to growth in cell culture conditions^53^. As a result, we were thus unable to explore the generalizability of our findings across different TNBC models from different genetic backgrounds. The reader should keep this fact in mind when interpreting our data. In future, a key priority will be to characterize a wider panel of models that recapitulate physiological binding between CD44 and HA. Patient-derived xenografts (PDX) or 3-D spheroid cultures are becoming more widely applied in breast cancer research and may be useful models for studying the connections between RAB4A, CD44 and HA binding.

In addition to our work on RAB4A, our CRISPR screen also provides a useful resource for discovery of other factors that may influence CD44-HA binding in breast cancer. We would refer to the reader to Fig. 2E for a curated list of possible targets of interest. While RAB4A stood out as being genetically amplified in many breast cancer patients, a review of the literature reveals links between several genes on this list and cancer metastasis. CLDN9, for example, is a cell membrane protein that is a component of tight junctions^54^. CLDN9 is a member of the claudin protein family, which are key regulators of cell migration and adhesion in cancer^55^. LAMC2, additionally, is a secreted glycoprotein that is thought to be a component of laminin in the basement membrane ECM^56^. In previous work, each of these proteins has been shown to physically interact with specific cell-surface receptors and regulate their localization to the plasma membrane^39,57-59^. Additionally, both have been mechanistically linked to tumor cell invasion both *in vitro* and *in vivo*^39,57,60^. No connection has ever been made, however, between these genes and regulation of CD44-HA binding. Such molecules are thus potentially high-value targets for modulation of the CD44-HA axis.

Most importantly, our results validate a general strategy for dissecting interactions between glycosaminoglycans (GAGs) and their cognate cell-surface receptors. Cancer cell invasion and migration is mediated by interactions with a variety of heavily glycosylated macromolecules in the ECM. Our FACS-based screening approach could easily be adapted to study the biology of other ECM glycopolymers such as heparin sulfate proteoglycans, chondroitin sulfate proteoglycans and/or mucin-type O-glycoproteins. Similarly, it would be straightforward to apply other CRISPR technologies, such as CRISPR activation and/or base editor screening, as a part of the same overall screening pipeline. We thus envision significant future opportunities to apply this technology to explore other aspects of GAG biology in cancer.

## Acknowledgements

S.W., P.J. and C.D.R. acknowledge funding from the Canadian Glycomics Network (GlycoNet) and the Cancer Research Society.

## Materials & Methods

### Cell Culture

MDA-MB-231 cells (acquired from the American Type Culture Collection) and Lenti-X cells (acquired from Takara) were cultured at low passages in Dulbecco’s Modified Eagle’s Medium (DMEM; Gibco, Thermo Fisher) supplemented with 10% Fetal Bovine Serum (FBS; Gibco) and maintained at 5% CO2 and 37°C. Trypsin-EDTA (0.25%, Gibco) and TrypLE (Gibco) were used to harvest cell cultures from tissue culture polystyrene.

### Lentiviral transduction of MDA-MB-231 cells

For lentiviral packaging, 7.5 x 10^6^ Lenti-X cells were seeded in a 150 mm tissue culture dish and transfected with 48 µL of TransIT-LT1 (Mirus, Cat # MIR2304), 8 µg of packaging plasmids (4 µg psPAX2, 4 µg pMD2.G) and 8 µg of the sgRNA-encoding plasmid. Lentiviral supernatant was harvested 72 hours after transfection and concentrated 10x using the Lenti-X concentrator reagent (Takara, Cat # 631231). For transduction, cells were transduced with concentrated lentiviral supernatant along with polybrene (8 µg/mL) for 48 hours. Cells were subsequently cultured with an appropriate selection antibiotic until the cells on a non-transfected control plate had been fully depleted. For CRISPR KO experiments, an sgRNA guide targeting RAB4A (sgRNA sequence: GTGACGAGAAGTTATTACCG) was cloned into a LentiCRISPR-v2 (Addgene, Cat # 52961) vector backbone, which encodes both the relevant sgRNA and Cas9, and selected using puromycin (1 µg/ml). For CRISPR KD experiments, MDA-MB-231 cells were first transduced with a lentivirus encoding dCas9KRAB (Cellecta, Cat # SVKRABC9B-PS) and selected using blasticidin (InvivoGen, Cat # ant-bl-05, 5 µg/mL). Cells were then transduced with a plasmid (AddGene Cat # 60955) encoding the sgRNA sequence GGCAGCCGCTGGGAGACCGG and selected with puromycin (InvivoGen, Cat # ant-pr-1, 1 µg/mL).

### Flow cytometry

1×10^5^ MDA-MB-231 WT, CD44KD, RAB4A KO and respective sgNT control cells were harvested and washed with PBS before being spun down at 600xg for 5 minutes. FL-HA was produced in-house according to a previously established protocol^22^. For FL-HA staining, cells were resuspended in FACS buffer (PBS + 2% FBS) containing a 1:100 dilution of FL-HA for 30 minutes on ice in a 96-well V-bottom. To also ensure FL-HA bound to CD44, WT cells were initially harvested in serum-free DMEM media and treated with a CD44 Hermes-1 blocker (1 µL per 10,000 cells) at 37°C for 1 hour. The blocker was then quenched by adding 950 µL of serum-free media. Cells were spun down, washed with PBS and stained with FL-HA. For CD44 staining, a 1:100 dilution of anti-CD44 [Hermes-3] (Abcam, Cat # 254530) was pre-complexed with 1.5 µg/mL goat anti-mouse Alexa 488 (Jackson ImmunoResearch, Cat # 115-545-071) in FACS buffer for 30 minutes on ice. The same was done with a mouse IgG2a isotype control (R&D, Cat # MAB003). Cells were subsequently centrifuged and washed with cold PBS to be analyzed by flow cytometry. They were transferred to a 96-well U-bottom plate and run on a benchtop Aurora flow cytometer (Cytek Biosciences). Gating on the forward/side scatter was used to identify live cells. The FITC-B2-A channel was used to gate FL-HA+ or CD44+ cells. Data was analyzed using the FlowJo software (Becton Dickinson, version 10.8.2)

### Genome-wide CRISPRi screening

MDA-dCas9–KRAB cells were transduced with a genome-wide CRISPRi library of sgRNAs consisting of 5 sgRNAs/gene (AddGene, Cat # 83969) as described above. They were subsequently stained with FL-HA as described above and sorted by FACS to isolate a “high-staining” (top 20% of the fluorescence distribution) and “low-staining” (bottom 20%) population of cells. Library amplification by PCR and Illumina sequencing was performed as described previously^61^.

### Bioinformatic analysis

CRISPR screen analysis was performed using MaGeCK^28^ according to a previously established protocol^61^. A single score was calculated for every gene by subtracting the -log_10_(negative selection score) from the -log_10_(positive selection score). This step arbitrarily assigns a positive value to hits that decreased FL-HA binding and a negative value to hits that increased FL-HA binding. TCGA analysis was conducted by accessing data associated with the cited cohort^34^ using cBioPortal. GO term analysis was performed using GOrilla^62^ and essentiality analysis was performed using DepMap^63^. For EnrichR analysis of differentially expressed proteins, all genes with log2(foldchange) less than -2 in RAB4A KO cells and an FDR of less than 0.01 were defined as a target list. The total list of proteins detected in the proteomics dataset were defined as the background list^64^. For EnrichR analysis of proteins downregulated at the protein but not the mRNA level, all genes with log2(foldchange) less than -2 in our proteomics dataset and greater than -1 in our transcriptomics dataset were defined as a target list. The total list of proteins detected in the proteomics dataset was defined as our background list. Comparison of RAB4A gene expression in normal, primary tumor and metastatic tumor samples was performed using TMNplot^65^. Survival analysis was performed with Kaplan-Meier plotter^66^. RFS was used as an endpoint for analysis of 1671 basal-type cancer samples. Cutoffs were autoselected. CD44/RAB4A mRNA expression correlation analysis was performed on basal-like samples within the METABRIC cohort using cBioPortal.

### DNA extraction and amplification

Genomic DNA was extracted from MDA-MB-231 WT and RAB4A KO cells using the GenElute™ Mammalian Genomic DNA Miniprep Kit (Sigma Life Science, Cat # G1N70) according to manufacturer’s instructions. Forward (CTATGGATTCCACCGCGCC) and reverse (CTGGCCTGAGAGGGTCAATC) primers were obtained from Integrated DNA Technologies (IDT). A Polymerase Chain Reaction (PCR) was performed in 50 µL reaction volumes using 300 ng DNA template, 5 µL of each primer (10 µM), 11 µL of 5X Herculase II Reaction Buffer, Herculase II Fusion DNA Polymerase Enzyme, 100 nM dNTP, and topped up with nuclease-free water. The Herculase II Enzyme with dNTP combo was obtained from Agilent (Cat No. 600679). The reactions were performed with 2 minutes of initial denaturation at 98ºC, followed by 35 cycles of 30 seconds of denaturation at 98ºC, 30 seconds of annealing at 55ºC, and 1 minute of extension at 72ºC. In the final extension step, the reaction was held at 72ºC for 3 minutes. PCR products were then run on a 1% agarose gel and the visible band was carefully excised. PCR products were extracted using the GeneJET Gel Extraction Kit (Thermo Scientific™, Cat # K0692).

### Sanger sequencing and knockout efficiency analysis

Purified PCR products from amplification of MDA-MD-231 WT and RAB4A KO cells were prepared and submitted for Sanger sequencing in accordance with UBC Sequencing and Bioinformatics Consortium pre-defined guidelines. Knockout efficiency was determined using the ICE CRISPR Analysis Tool by Synthego (https://ice.editco.bio/#/).

### Western blot

The efficiency of RAB4A KO was confirmed using Western blots. Pierce® RIPA Buffer (Thermo Fisher Scientific, Cat # 8990) with Halt™ Protease Inhibitors (Thermo Scientific, Cat # 78430) were used to lyse MDA-MB-231 WT and KO cells. The supernatant was collected, and protein concentration was determined using the Pierce™ BCA Protein Assay (Cat # 55864) according to manufacturer’s instructions. Lysate consisting of 20 µg protein was heated to 95ºC and run in each lane of 10-well, 4–15% Mini-PROTEAN TGX Precast Protein Gel (BioRad, Cat # 4561083). Separated protein was transferred using the Trans-blot® Turbo Transfer System (Bio-Rad Laboratories) to midi-size 0.2 um nitrocellulose transfer membrane (BioRad, Cat # 1704159). The membrane was blocked with blocking buffer consisting of 5% BSA in PBS containing 0.5% Tween-20 (BSA: Sigma-Aldrich, Product No. 81053; Tween 20: Thermo Fisher; Cat No. J20605.AP) at room temperature for 60 minutes. Primary recombinant mouse monoclonal anti-CD44 antibody (Abcam, Ab254530, 1:1000 dilution), rabbit monoclonal anti-RAB4A antibody (Abcam, EPR3042, 1:5000 dilution), and anti-GAPDH antibody (Cell Signaling Technology; Cat No. 97166S 1:1000 dilution) were resuspended in TBS with 0.1% Tween20 (TBS-T) and incubated with the membranes in a cold room overnight. Goat anti-Rabbit (IRDye® 800CW; LICORbio) and Goat anti-Mouse (IRDye® 680RD; LICORbio,) were diluted 1:10000 in the blocking buffer and used as secondary antibodies for anti-RAB4A and anti-CD44/anti-GAPDH respectively. The membranes were incubated with secondary antibodies at room temperature for 60 minutes, then washed with TBS-T and imaged using the Sapphire Biomolecular Imager (Azure Biosystems).

### Immunofluorescence

Ibidi 8-well chamber slides (Ibidi, Cat # 80826) were treated with 150 μL of Poly-D-Lysine (Gibco, Cat # A38904-01) for 1 hour at room temperature. Excess Poly-D-Lysine was removed, and the chamber slide was washed twice with sterile PBS. 35,000 WT, sgCD44 and sgRAB4A cells were seeded and allowed to rest overnight at 5% CO2 and 37°C. The next day, cells were fixed with cold 4% PFA for 15 minutes. They were washed thrice with 150 µL PBS for 5 minutes each to remove excess PFA. The cells were then permeabilized with 0.3% Triton in PBS for 15 minutes and blocked with 1% BSA in PBS for an hour at room temperature. Cells were then incubated with a primary antibody against CD44 (Abcam, Cat # 254530) at a 1:500 dilution overnight at 4°C. The next day, cells were incubated with fluorescent secondaries using anti-mouse Alexa 488 (Jackson ImmunoResearch, Cat # 115-545-071) for one hour at room temperature on a rocker. Post-secondary incubation, 2 washes with 1:1500 diluted DAPI (Invitrogen, Cat # 62248) were done for 5 minutes each. Between each step, cells were washed with PBS three times for 5 minutes each on a gentle rocker. Cells were then kept in PBS and imaged using an LSM900 confocal microscope. Laser powers for all channels were set at 0.8% and the gain at 750V.

### Transwell invasion assay

Transparent PET membrane inserts with 8 μm pores (Falcon®, 353097) were coated with 50 μL of 1 mg/mL high-molecular-weight HA (Sigma, H5388-100MG) and placed into a 24-well plate. The plates with transwells were left at room temperature overnight to allow the formation of solidified HA-containing matrices. MDA-MB-231 sgNT, CD44 KD and RAB4A KO cells were harvested in serum-free DMEM the next day. Before adding the cells, 500 μL of DMEM supplemented with 10% FBS were added to the bottom of the 24-well plate below the transwells to form a gradient. 50,000 cells in serum-free media were seeded onto the top of the transwells such that the total volume was maintained at 100 μL. The transwells were incubated at 5% CO2 and 37°C for 48 hours. Post incubation, uninvaded cells and remaining HA matrix were scraped off with a dry Q-tip once and washed twice with Q-tips dipped in PBS. Invaded cells thus remained at the bottom of the transwells. The wells were fixed in 100% methanol for 15 minutes and stained with 0.5% crystal violet in 80% methanol for 20 minutes. The wells were then dipped into distilled water as washes to remove excess crystal violet. After allowing the crystal violet to dry, cells on each well were visualized on a bench-top phase contrast microscope and counted using 3×3 gridded 60mm dishes. Number of invaded cells were counted across the entire 9 fields.

### Real Time-Quantitative PCR (RT-qPCR) analysis of CD44 splice variants

Primers to screen for expression of variants within a part of the human CD44 gene were obtained from a previously described protocol^67^ and ordered from IDT. A reverse primer, C2A, served as a common primer for all forward primers located on different exons of the CD44 gene. Primers are provided below. The total RNA from sgNT and sgRAB4A cells were extracted (Monarch® Total RNA Miniprep Kit) and the corresponding cDNA was synthesized using the protocols provided by the manufacturer (LunaScript® RT SuperMix Kit). The cDNA from each sample was added to a pre-made qPCR mastermix (Luna® Universal qPCR Master Mix) along with the forward and common reverse primers. Each sample was run using the standard two-hour protocol on the StepOnePlus System (Applied Biosystems), and the Ct values obtained were analyzed using Microsoft Excel.

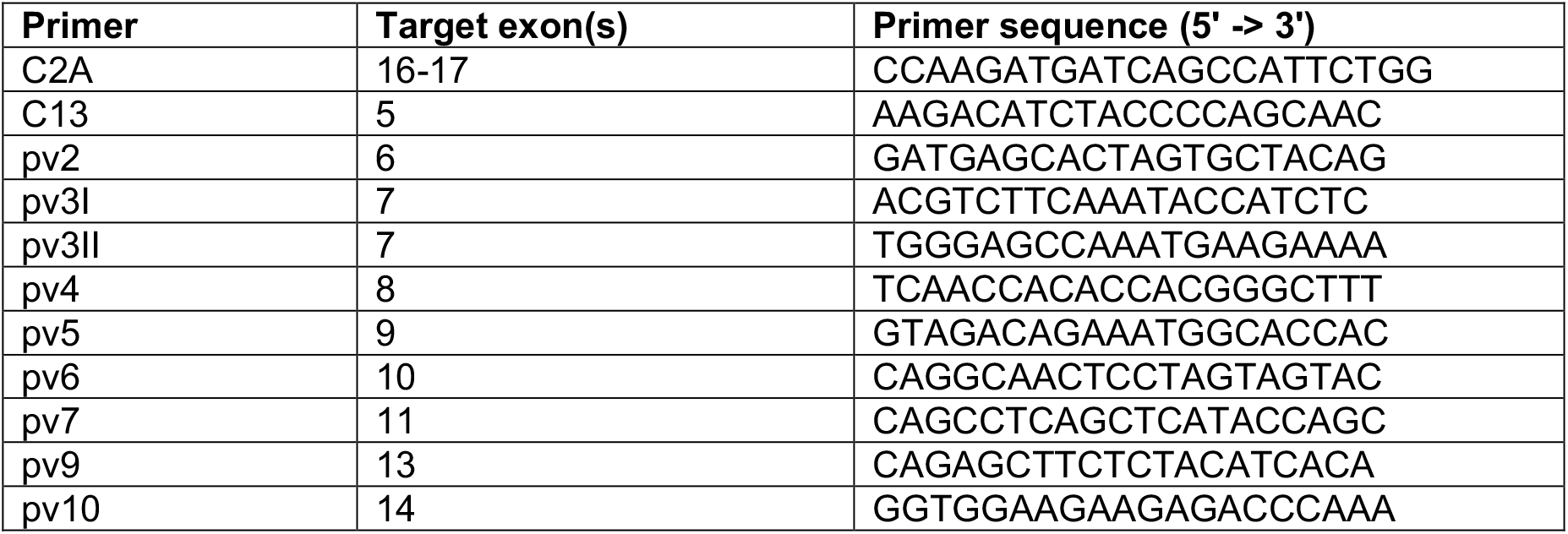

For analysis of RAB4A expression in WT and RAB4A KD cells, RT-qPCR was performed as above using the primers CGCTCCCAAGATGTCGCAGA (Forward) and ACTGATGAAGTAAGCAAGATTTGCC (Reverse).

### Statistical analysis

In all flow cytometry and qPCR experiments, a standard Student’s unpaired *t*-test was applied as a standard method for determining statistical significance. For transwells, a One-Way ANOVA was applied for determining statistical significance. Further statistical details are included in figure legends. No statistical method was used to determine sample size. Controls were included as appropriate to provide a reference with which to compare experimental data. No data were excluded from any analyses. All data were confirmed with multiple biological replicates as indicated in the figure legends.

